# Identifying associations of *de novo* noncoding variants with autism through integration of gene expression, sequence and sex information

**DOI:** 10.1101/2024.03.20.585624

**Authors:** Runjia Li, Jason Ernst

**Author notes:** Correspondence (JE).

## Abstract

Whole-genome sequencing (WGS) data is facilitating genome-wide identification of rare noncoding variants, while elucidating their roles in disease remains challenging. Towards this end, we first revisit a reported significant brain-related association signal of autism spectrum disorder (ASD) detected from *de novo* noncoding variants attributed to deep-learning and show that local GC content can capture similar association signals. We further show that the association signal appears driven by variants from male proband-female sibling pairs that are upstream of assigned genes. We then develop Expression Neighborhood Sequence Association Study (ENSAS), which utilizes gene expression correlations and sequence information, to more systematically identify phenotype-associated variant sets. Applying ENSAS to the same set of *de novo* variants, we identify gene expression-based neighborhoods showing significant ASD association signal, enriched for synapse-related gene ontology terms. For these top neighborhoods, we also identify chromatin states annotations of variants that are predictive of the proband-sibling local GC content differences. Our work provides new insights into associations of non-coding *de novo* mutations in ASD and presents an analytical framework applicable to other phenotypes.

## Introduction

The increasing availability of whole genome sequencing (WGS) data is presenting new opportunities to better understand the associations of rare noncoding variants with complex human phenotypes^1–3^. One such phenotype where WGS has been applied is to study genetic contributions to autism spectrum disorder (ASD). Prior studies using arrays or exome sequencing led to genetic associations with ASD based on common^4^ or rare coding^5–10^ variants, respectively, but were not able to extensively study the role of rare noncoding variants. A notable dataset that has been used to begin to understand the role of rare noncoding variants in ASD and more broadly psychiatric and other complex diseases is WGS of the Simons Simplex Collection (SSC) cohort^3,11–14^. This dataset has enabled calling of *de novo* noncoding variants from WGS of probands and unaffected siblings for more than a thousand families. *De novo* variants are of particular interest^15^ since they correspond to a subset of variants for which selection has not had a chance to exert its effect and thus can enrich for higher impact variants^16^.

However even when focusing on *de novo* noncoding variants, there remain substantial analytic challenges in interpreting the association of these variants to ASD. In an initial analysis of a subset of the *de novo* variants in the SSC cohort, many combinations of variant annotations were tested for burden association through the category-wide association study (CWAS) framework^12^. After multiple-testing correction, this did not lead to any significant noncoding associations in either an initial cohort^12^ or later analysis of an expanded cohort^13^. However, an alternative analysis based on a risk score did suggest a significant noncoding association signal and further analysis suggested that this signal was associated with promoter regions^13^.

A separate analysis of the SSC cohort defined a DNA-based and an RNA-based ‘disease impact score’ (DIS), which were then used to evaluate the contributions of *de novo* noncoding variants to ASD^11^. The scores were generated based on first training deep neural networks to predict from DNA sequences chromatin accessibility, histone modifications, transcription factor (TF) binding, and RNA binding. The trained neural networks were then used to compute corresponding feature values for individual variants based on their impact on the neural network predictions. Finally, DNA- and RNA-based features were combined into their respective DISs with a supervised classifier trained using disease-curated variants. In addition, a DIS that combines the DNA- and RNA-based DIS scores was also defined. Various subsets of de novo variants from the SSC cohort were then tested to determine if there was a significant difference in DIS distributions of them between proband and unaffected siblings. This led to reported significant associations including notably among a subset of noncoding variants that are near genes that are differentially expressed in brain tissues.

However, given the complexity of the approach and the challenge of interpreting a score based on integrating many different features derived from deep neural networks, we sought to investigate whether simpler and more interpretable approaches could identify a similar or even stronger association signal. Specifically, here we first show that using local GC content is sufficient to obtain similar results about noncoding variant association previously attributed to deep learning. We then further show that by considering simple additional information not considered in the prior analyses, namely the sex of the proband and the unaffected sibling and whether the variant was upstream or downstream of their assigned genes, we can better isolate the likely source of the association signal for variants near genes differentially expressed in brain. Specifically, we found that the association signal for variants near differentially expressed brain genes appears to be driven by variants from families with a male proband and female sibling and are upstream of the gene they are assigned.

Given these insights, we develop Expression Neighborhood Sequence Association Study (ENSAS) to extend the analytical approach to also evaluate high-order k-mers and more comprehensively defined gene-expression variant sets. ENSAS identifies sets of variants associated with a phenotype by integrating systematically defined gene expression neighborhoods and sequence context in the form of k-mers. We apply ENSAS to analyze the SSC cohort with ASD. When considering variants from male probands and female siblings upstream of their assigned genes, ENSAS refines the set of brain-expressed gene sets most strongly associated with the association signal. Gene ontology enrichment analyses suggest the top associated neighborhoods are related to synapses. Using higher-order k-mers either does not improve predictive performance or shows at most a limited improvement over local GC content. Finally, we conduct chromatin state analyses of variants in top neighborhoods which suggest that chromatin state annotations from specific epigenomes can predict a substantial portion of the proband-sibling GC content differences. Together our work provides new insights into *de novo* noncoding mutations associations with ASD and presents a general analytical framework that could be used for other phenotypes.

## Results

### Local GC content largely explains noncoding ASD associations attributed to deep learning

We first investigated if a simpler and more interpretable procedure can yield similar association signals compared to using DIS score. In particular, across the n=127,140 *de novo* variants we observed a strong correlation between local GC content, defined here as the number of G or C bases in the 201bp window centered at the variant, and both the RNA DIS and the DNA DIS (Spearman correlation=0.57 and 0.72, respectively) (Figure 1). This led us to ask whether similar association signals could be obtained with just local GC content.

**Figure 1.**
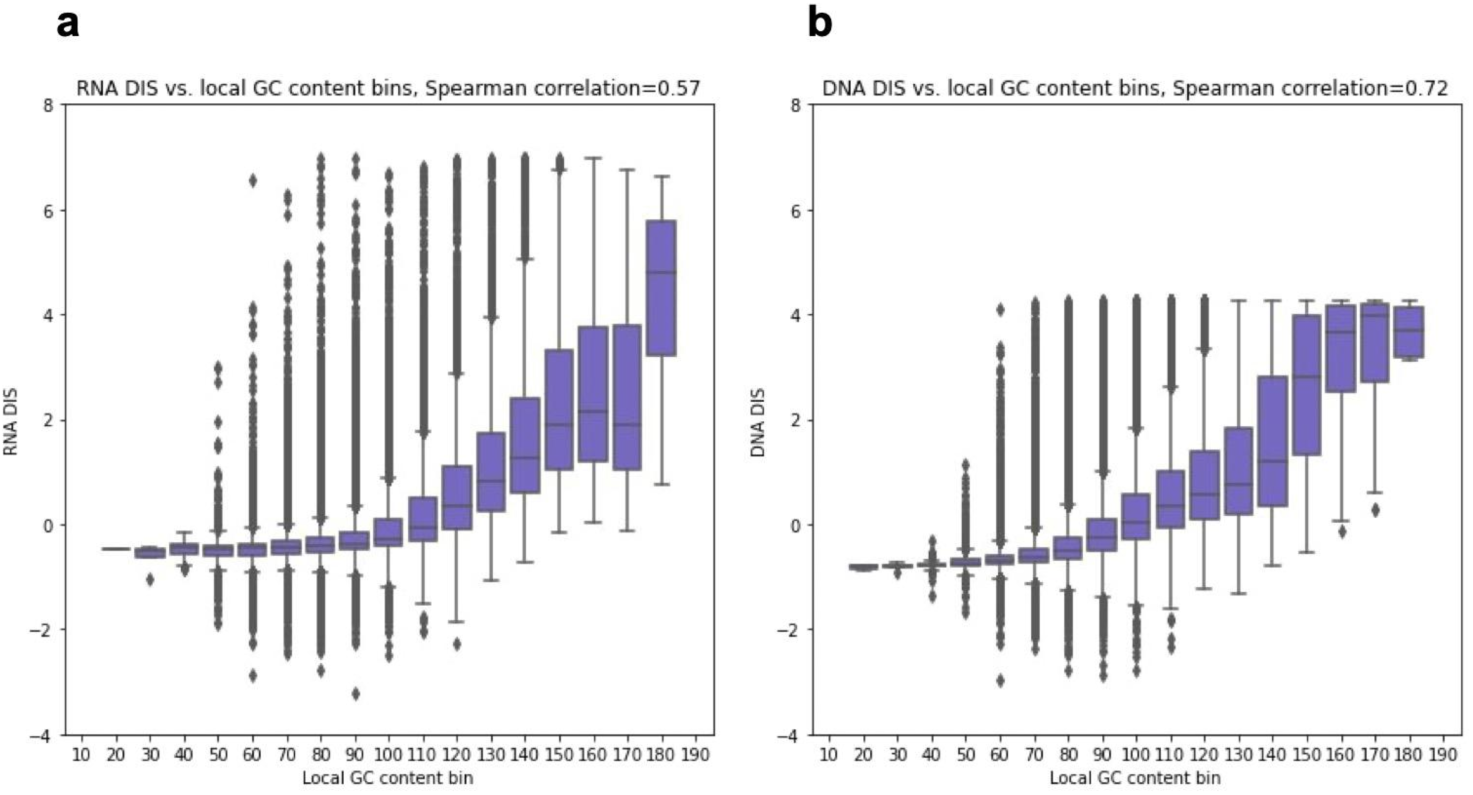
DIS vs. local GC content. **(a,b)** The y-axis corresponds to a DIS score and the x-axis 20 evenly sized local GC content bins ranging from 0 to 201. Each bin is represented by a box showing the distribution of its variant’s DIS^11^. The boxes correspond to the quartiles, the lengths of whiskers correspond to 1.5x interquartile range and variants above/below the whiskers are defined as outliers. Spearman correlations are computed between DIS and local GC content, across all variants. **(a)** RNA DIS. **(b)** DNA DIS.

To test this we conducted comparisons using the same association tests previously reported for variant sets defined for (1) 130 curated genomic variant sets (60 for DNA DIS and 70 for RNA DIS) (Supplementary Figure 1) and (2) proximity to genes exhibiting tissue-restricted expression activity based on expression data in 53 tissues or cell types from the Genotype–Tissue Expression (GTEx) project^17^ (Figure 2a, b). Here for the genomic variant set analysis we used local GC content in place of the RNA or DNA DIS and for the GTEx-based analysis in place of the combined DIS. For the GTEx-based analysis following Zhou *et al*.^11^ we only included variants within 100kbp of a transcription start site (TSS) or intron variants within 400bp of an exon boundary, which left 71,554 variants. We also conducted additional tests where we excluded both coding and canonical splice site (CSS) variants from the analysis, both of which have well-established associations with ASD^4,5,7–10,14^, which left us with 122,807 variants for the genomic variant set analysis and 67,671 variants for the GTEx-based analysis.

**Figure 2.**
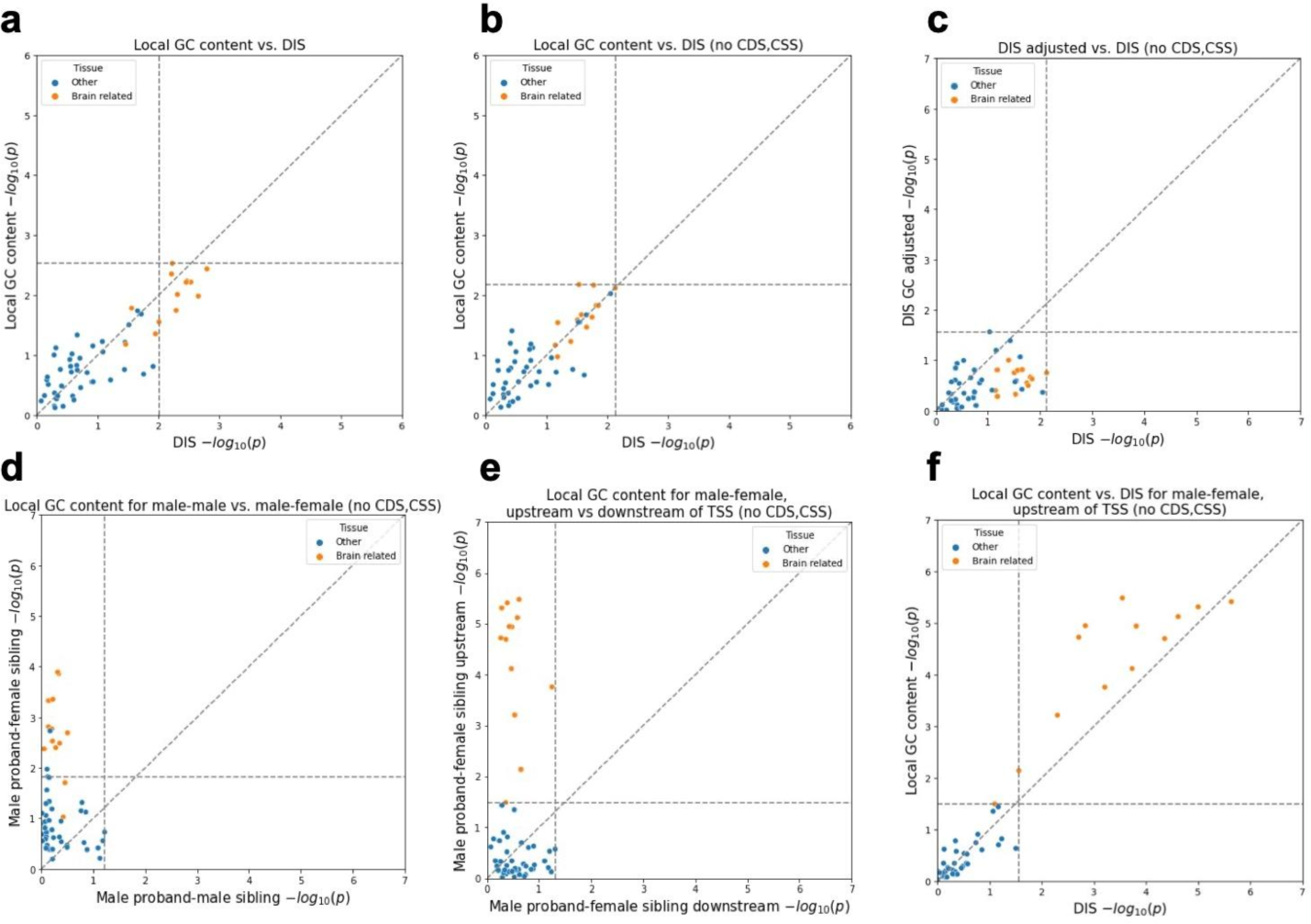
Proband-sibling differences for DIS (combined) or local GC content of variants assigned to each of the 53 GTEx tissue or cell types. Both the x- and y-axis show -log10 p-values from one-sided Mann-Whitney U-tests. Horizontal and vertical dashed lines show p-values at FDR threshold of 0.05. Points greater than (but not on) these lines are significant after FDR correction. Diagonal dashed lines show the unit slope. **(a)** Proband-sibling differences for proband-sibling differences for DIS (as conducted by Zhou *et al*.^11^) vs. local GC content; **(b)** same as (a) but with coding and CSS variants removed; **(c)** proband-sibling differences for DIS vs. proband-sibling differences for DIS (adjusted for local GC content), with coding and CSS variants removed; **(d)** male proband-female sibling pair differences for local GC content vs. male proband-male sibling pair differences for local GC content, with coding and CSS variants removed; **(e)** male proband-female sibling pair differences for local GC content in variants <100kbp upstream of nearest outermost TSS only vs. in variants <100kbp downstream of nearest outermost TSS, with coding and CSS variants removed. **(f)** similar to (a) but restricted to male proband-female sibling pair variants < 100kbp upstream of nearest outermost TSS, with coding and CSS variants removed.The specific values corresponding to this analysis can be found in Supplementary Table 1.

Using local GC content we obtained an association signal with an overall similar distribution of p-values for the DNA DIS across the 60 curated variant sets (two-sided Mann-Whitney U p=0.16) and the combined DIS across the 53 GTEx tissues (p=0.98). The RNA DIS showed stronger associations compared to local GC content when considering the 70 variant sets based on all variants (p=0.009; Supplementary Figure 1a, b), but after excluding coding and CSS variants the difference was no longer significant (p=0.16, Supplementary Figure 1c, d). To more directly test whether the reported DIS significant associations signal could be explained by local GC content we adjusted the DIS score based on the local GC content (Methods). After adjusting for local GC content and controlling for multiple testing individual associations were no longer significant (Figure 2c, Supplementary Figure 1e, f).

While these analyses suggest local GC content offers a simpler approach sufficient to produce similar association signals to those reported for DIS, they do not exclude the possibility that there is additional association signal within the DNA sequence beyond local GC content. These analyses also do not explain why local GC content, which can be both biologically significant and a confounder in genomic analyses^18,19^, was sufficient to identify the association signals. In particular, we were interested to understand this in the context of the result that variants assigned to genes differentially expressed in brain tissue types among a panel of GTEx tissue types had the strongest association signal for ASD.

### Local GC content differences around brain-specific genes are specific to male probands with female siblings

Given the well-established sex bias of ASD cases^20^, which was reflected in the SSC cohort with 87% of probands male compared to 47% of unaffected siblings, we asked if sex differences between probands and siblings might be related to the local GC content differences between their *de novo* variants assigned to brain-specific expressed genes. Specifically, we repeated the gene expression association tests based on local GC content separately for *de novo* variants from a family with a male proband and a female sibling (n=30,430) and for variants from a family with a male proband and a male sibling (n=27,251). We note that consistent with the previous analysis all variants that we considered were on autosomes and excluded those variants overlapping repeats^11^. Variants from male proband-female sibling pairs showed noticeably stronger signals than those from male proband-male sibling pairs. For male proband-female sibling pairs 13 tissues were significant at an FDR threshold of 0.05 with the most significant tissue having a nominal p-value of 0.00013 compared to no tissues significant at the same FDR threshold for male proband-male sibling pairs and the most significant tissue having a nominal p-value of 0.061 (Figure 2d). Brain-related tissues were among the top associations for male proband-female sibling pairs with 11 out of 13 tissues significant at an FDR threshold of 0.05 but not for male proband-male sibling pairs where the most significant brain-related tissue had a p-value of 0.31 (Figure 2d). These results suggest that the brain-related signal is mainly associated with male proband-female sibling pairs.

### Male proband-female sibling differences in local GC content are specific to variants upstream of TSSs

In the above analyses, we used the previous assignments of variants to genes based on their nearest representative TSS^11^, which did not differentiate variants upstream of a TSS from those downstream. Given the different chromatin environments and mutational processes associated with transcription^21^, we reasoned that it could also be informative to analyze the association signal separately for variants upstream and downstream of the TSS of their assigned genes. Specifically, we computed for each variant its position relative to the nearest annotated outermost TSS of a protein-coding gene, restricting to variants within 100kbp of a TSS, consistent with the distance threshold previously used^11^. We then repeated the above gene expression-based analysis separately for variants from male proband-female sibling pairs that were upstream of their assigned gene and for those downstream. Overall, when restricted to the set of upstream variants (n=12,474) the brain tissues showed even stronger association signals compared to using all variants, with 12 tissues significant at an FDR threshold of 0.05 and the most significant tissue being the anterior cingulate cortex (nominal p=3.3×10^-6^). Meanwhile none of the tissues are significant for the downstream variants (n=16,571), with the most significant tissue being skeletal muscle (nominal p=0.049) and the most significant brain-related tissue being hypothalamus (nominal p=0.057). Compared to using the combined DIS for this analysis, local GC content showed an overall similar trend but had a more significant p-value for 12 of the 13 brain tissues (Figure 2f). Together, these results suggest the observed difference in local GC content between male proband and female sibling variants near brain-expressed genes can be mainly attributed to variants upstream of TSS.

### Analyses of the brain-related signal with Expression Neighborhood Sequence Association Study

The previous gene expression analyses were limited to using previously defined sets of genes for each of the 53 GTEx tissues, with each set being the genes that in the tissue type had five times its median expression across all tissues. This leaves open the possibility that considering information from additional gene expression data and more comprehensively defined gene expression based gene sets could capture potential additional signals. Additionally, the previous analyses were limited to only using local GC content in terms of sequence information, leaving open the possibility there is additional association signal information in the sequences that could be captured by higher order k-mers.

We therefore propose Expression Neighborhood Sequence Association Study (ENSAS, Methods), a generalized framework that integrates gene expression neighborhoods and local sequence in the form of k-mers counts to discover sets of variants associated with a phenotype (Figure 3). ENSAS first defines gene-expression-based neighborhoods by considering pairwise gene expression correlations. Specifically, ENSAS uses previously computed gene-gene expression correlations from the Geneshot database^22^, which were computed using a large compendium of gene expression datasets collected by the ARCHS4 web resource^23^. Compared to GTEx, ARCHS4 is a more comprehensive and diverse collection of publicly available RNA-seq data, consisting of over 80,000 human samples across different tissues^23^. ENSAS then defines a variant ‘neighborhood’ surrounding each gene, as the variants assigned to the gene itself and the top M variants assigned to a set of closest genes based on the expression correlations (Methods).

**Figure 3.**
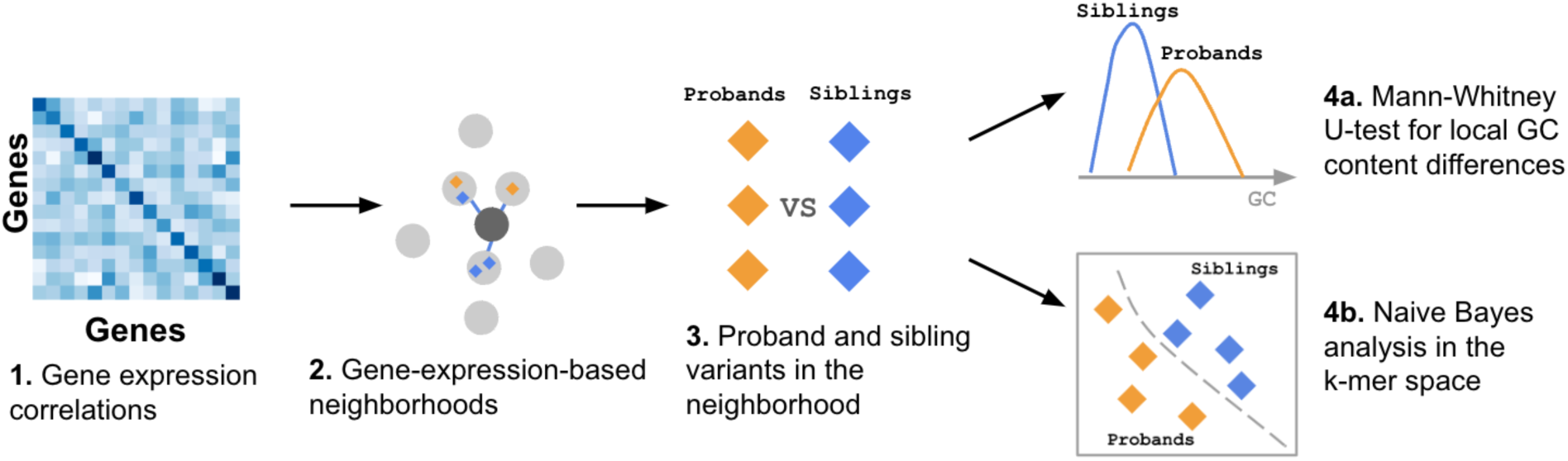
Schematic of the ENSAS. In this example, the two sets of variants being compared are the proband vs.unaffected sibling variants. **(1)** The gene-gene expression correlation matrix is obtained from the Geneshot database^22^. **(2)** ENSAS identifies the genes with the largest gene expression correlations to a target gene and the neighborhood of the target gene is defined as the top M variants assigned to its closest genes based on the correlations, including the variants assigned to the target gene itself if there are any. **(3)** Each neighborhood consists of two groups of variants which are compared using the following approaches: **(4)** ENSAS first (a) performs a Mann-Whitney U test between the local GC content of proband and sibling variants, and additionally (b) uses a Naive Bayes model to identify sequence differences based on k-mers.

After defining gene-expression-based neighborhoods, ENSAS conducts two sets of sequence based tests for each neighborhood. The first is a one-sided Mann-Whitney U-test to test the difference between the local GC content of two groups of variants. The second is a test based on k-mer frequencies. Specifically, for this second test ENSAS splits the variants in a neighborhood into equally sized training and testing folds. Using variants from the training fold ENSAS trains a multinomial Naive Bayes classifier with uniform class priors to predict a variant’s group label. The features to the classifier are the k-mer counts within a L-bp sequence window centered on the variant. We used the Naive Bayes classifier as it is suitable for small sample sizes, and similar formulations of k-mer-based Naive Bayes classifiers have previously been used for sequence classification tasks^24–26^. ENSAS then uses the trained classifier to score each variant in the testing fold, where each score represents the variant’s relative likelihood of being in a target group. ENSAS then performs a one-sided Mann-Whitney U-test to test if the target group has higher scores than the non-target group. For comparison with k-mer results, ENSAS also conducts the local GC content analysis on the testing fold only.

Overall ENSAS can be seen as an extension of the previous GC content analysis with GTEx neighborhood, where an integration of k-mer frequency is tested in addition to GC content and the gene sets are neighborhoods surrounding each gene defined by gene expression correlations.

### ENSAS captures proband-sibling local GC content differences in specific gene-expression-based neighborhoods

We applied ENSAS to the SSC cohort, where the two groups are proband and sibling variants. For this application, we set the sequence length L to 201bp, the neighborhood sizes M to 1000 variants (mirroring approximately the mean GTEx variant set sizes that is 1048, Methods) and we used 1-7 as the candidate lengths of k-mers. We applied ENSAS restricted to variants from male-proband female sibling pairs that were within 100kbp upstream of their nearest outermost TSSs (henceforth referred to as “M-F upstream” variants) considered above, leaving us with n=12,293 variants. We tested a total of 29,820 expression neighborhoods and after a Bonferroni-based multiple testing correction we observed 28 neighborhoods with a significant difference (p<=1.7×10^-6^) in local GC content between proband and siblings. The top neighborhood consists of variants assigned to the closest genes surrounding OPCML, a synaptic signaling gene^27^ with a p-value of 1.3×10^-7^(Figure 4a). We note that the Bonferroni-based correction is conservative in this setting due to the overlap between neighborhoods, and ENSAS offers an alternative permutation-based multiple testing correction method though with a greater computational cost. Using a permutation based threshold (p<=2.8×10^-3^) we observed 1,542 significant neighborhoods when applied to the M-F upstream variants (Figure 4a, Methods). In the interest of analyzing a more limited set and the most significant neighborhoods, we focused our analyses on the 28 neighborhoods that were still significant after applying a Bonferroni correction.

**Figure 4.**
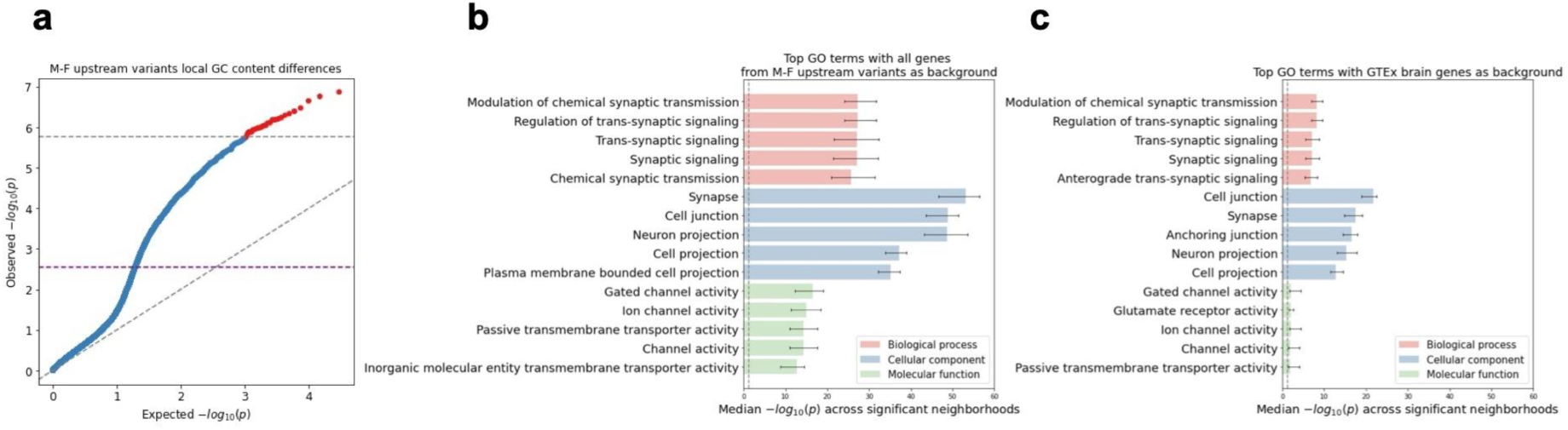
ENSAS results for M-F upstream variants in the SSC cohort. **(a)** QQ-plot of the neighborhood Mann-Whitney U p-values for local GC content. Grey horizontal dashed line shows the Bonferroni-based p-value multiple testing significance threshold of 0.05 / n where n=29,820 is the number of neighborhoods. Purple horizontal dashed line shows the permutation-based multiple testing threshold at an FDR of 0.05, with points above these lines significant after correction. Diagonal dashed line shows the unit slope. **(b)** Top 5 enriched GO terms under each GO category for the 28 significant neighborhoods using Bonferroni threshold (shown as red in (a)), using the union of all genes assigned to the M-F upstream variants as background. x-axis shows Benjamini-Hochberg adjusted Fisher’s exact p-values. Vertical dashed line shows p-value significance threshold of 0.05. Error bars show interquartile range across the neighborhoods. **(c)** Same as **(b)** but with the union of genes assigned to the M-F upstream variants that are also differentially expressed within any of the 13 brain-related GTEx tissues as background.

To biologically characterize the sets of genes associated with the top neighborhoods in the previous M-F upstream variants analyses we conducted a Gene Ontology (GO) enrichment analysis for genes associated with each of the 28 neighborhoods (Figure 4a, Methods). For this analysis we used as background the set of all genes assigned to M-F upstream variants. We observed that the most significant term across the neighborhoods was synapse (median Benjamini-Hochberg adjusted p=5.8×10^-54^, Figure 4b). We note the p-value significance in this analysis is the enrichment of genes in the top neighborhood for the GO category and not the significance of proband-sibling differences for the category. Other terms related to synaptic transmission and transmembrane channels were also highly significant. These classes of genes had previously been implicated based on *de novo* coding variants in ASD^6,28^.

We next asked whether the GO enrichments are more specific in the top neighborhoods identified by ENSAS or would be expected by also simply considering the set of previously analyzed differentially expressed genes in the 13 brain-related GTEx tissues. To investigate this we repeated the GO enrichment analysis but now using the set of genes assigned to the M-F variants that are within any of the 13 brain-related GTEx tissues as background. We observed that the top terms are similar as in the previous analysis, with the previously most significant term synapse still highly significant (median Benjamini-Hochberg adjusted p=3.5×10^-18^, Figure 4c). The most significant term is now cell junction (median Benjamini-Hochberg adjusted p=1.8×10^-22^, Figure 4c), which was previously also highly significant (p=1.2×10^-49^). That the top terms are preserved suggests the possibility that ENSAS identified top neighborhoods may provide more specific biological information compared to the union of differentially expressed genes in the 13 brain-related GTEx tissues.

### ENSAS Proband-sibling local GC content differences not driven by obvious sequencing batch effects or promoter variants

To exclude the possibility that the observed associations are driven by obvious sequencing batch effect as reflected in different recorded sequencing lanes for probands and unaffected siblings, we separately analyzed two subgroups of samples: samples from pairs that had matching recorded sequencing lane information and those from pairs that did not (Methods). This left us with 968 samples from male proband-female sibling pairs that had matching recorded sequencing lane information and 606 samples from pairs that did not. For each neighborhood, we separately performed proband vs. sibling Mann-Whitney U-tests on the two subgroups. When restricted to the samples with matching sequencing lane information, despite the smaller sample size, six significant neighborhoods were still identified as significant, with two of them also significant when considering the full set of samples (Supplementary Figure 2b).

The neighborhoods significant in the full set of samples had a median p-value of 1.7×10^-5^ in this subset of samples. No significant neighborhoods were identified when restricted to the samples with mismatched sequencing lanes (Supplementary Figure 2c) and the overall distribution of the p-values appeared relatively uniform (Supplementary Figure 2a). This suggests that the most of the signal can be attributed to a subset of samples whose comparison would be expected to be less susceptible to technical sequencing confounders.

We also confirmed that the signal identified here is distinct from the signal associated with variants in promoter regions in a previous analysis of the SSC cohort^13^, as would be expected as most variants we considered are outside of the promoter region. Specifically, from the initial set of 12,293 M-F upstream variants we removed those within 2kbp upstream of its assigned TSS following the previously used promoter definition^13^, which left 11,395 variants. We then applied ENSAS and observed a similar number of significant neighborhoods before and after removing promoter variants (28 vs. 21 neighborhoods, with 12 shared neighborhoods), with a Spearman correlation of 0.92 between the two sets of -log_10_ p-values (Supplementary Figure 3).

### Investigating sequence signal beyond local GC content in the significant neighborhoods

To first establish that ENSAS is powered to capture sequence context beyond local GC content, we simulated datasets containing variants from two groups with differences in their chromatin state distribution (Methods). We applied ENSAS to the simulated datasets and compared predictive performance using local GC content and Naive Bayes models with different k-mer lengths. We observed that under different simulation parameters (Methods) the best Naive Bayes models can consistently outperform local GC content (Supplementary Figure 4). These results confirm that a k-mer-based Naive Bayes model is capable of identifying association signals beyond local GC content.

We next applied ENSAS’s Naive Bayes analysis to investigate if the sequence context represented by k-mers is more predictive of proband sibling status compared to local GC content in specific neighborhoods. We separately used k-mers of lengths 1 to 7 as features and for comparison purposes local GC content was also evaluated on the same testing folds. None of the k-mer lengths led to improved overall performance of the Naive Bayes model over local GC content (Figure 5a). When restricted to the 28 neighborhoods with significant proband-sibling local GC content differences previously identified using all variants, the 6-mer model showed higher median performance compared to local GC content, but the difference did not reach statistical significance (one-sided Wilcoxon signed-rank p=0.34 for 6-mer, Figure 5b). When the random train-test splitting was repeated 100 times for each of these 28 neighborhoods, the 6-mer Naive Bayes model had better median performance in 18 of these neighborhoods (Supplementary Figure 5). Overall we did not see any k-mer Naive Bayes model significantly outperform local GC content in terms of proband-sibling association signal, but also cannot exclude the possibility that there exists additional sequence association signal beyond local GC content through 6-mers or other sequence representations.

**Figure 5.**
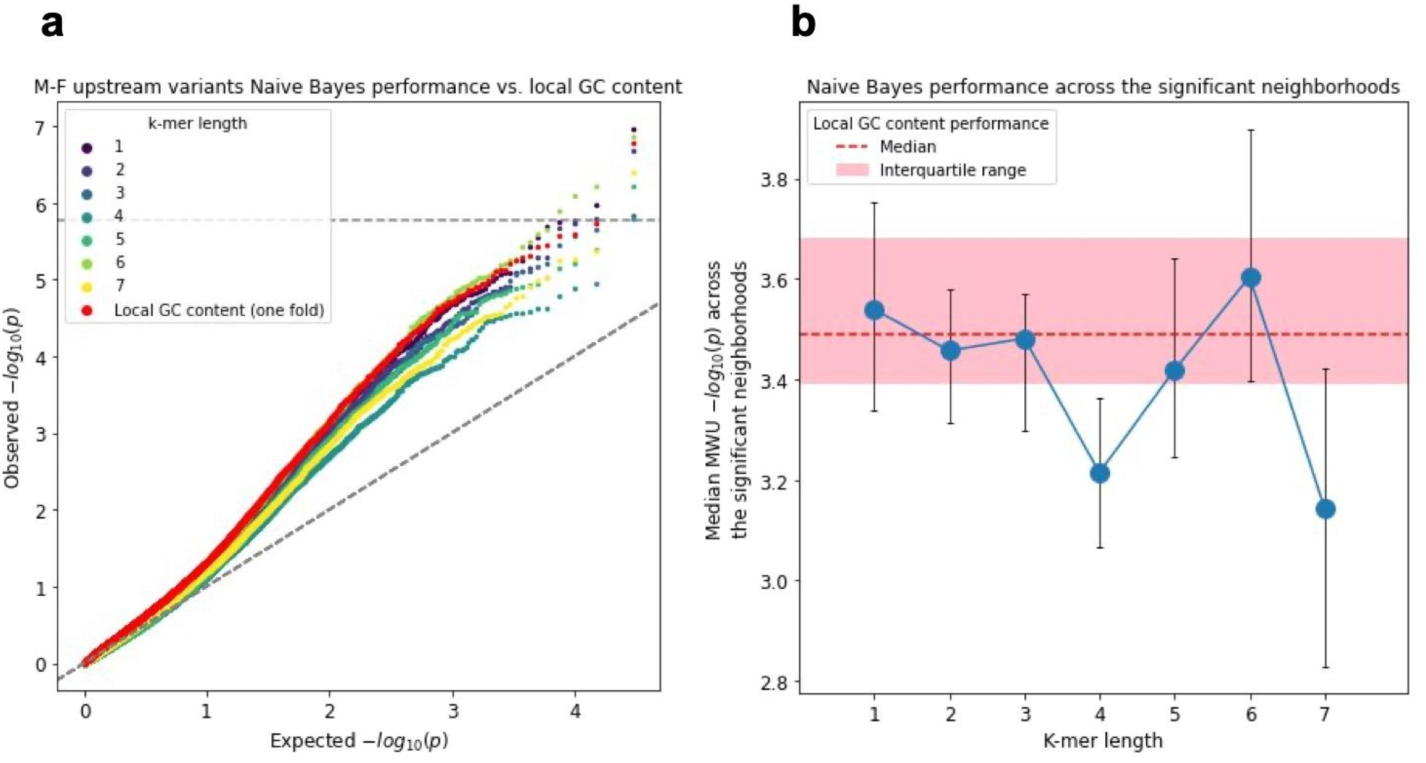
Naive Bayes model vs. local GC content predictive performance for the M-F upstream variant neighborhoods. **(a)** QQ-plot of the proband vs. sibling Mann-Whitney U p-values for the Naive Bayes scores, using k-mers of length 1-7 along with local GC content, all evaluated on the same testing folds. Diagonal dashed line shows the unit slope. **(b)** Median Mann-Whitney U p-values across the 28 significant neighborhoods identified based on local GC content using all variants for Naive Bayes models using k-mers of length 1-7, compared the local GC content (median p-value shown as red dashed line, interquartile range shown as shaded area). Error bars indicate interquartile range.

### Chromatin state annotation of variants in top neighborhoods is partially predictive of proband-sibling local GC content differences

We next investigated the extent to which the proband-sibling local GC content differences in the top neighborhood (surrounding OPCML gene) in the M-F upstream variants analysis is expected based on differences in the cell type-specific chromatin state annotations of the variants. For this we utilized annotations from a ChromHMM 18-chromatin state model defined based on six histone modifications that were available for 98-reference epigenomes from the Roadmap Epigenomics Consortium^29,30^. For each reference epigenome, we evaluated the extent to which the differences in local GC content ranks between proband and sibling variants are predicted by the variants’ chromatin state annotations (Methods). Across all epigenomes, the chromatin state annotations in an individual epigenome alone predicted a median of 19.1% of the local GC content differences, but reached as high as 50.2% predicted based on the annotations in a foreskin fibroblast sample. Among the epigenomes in the top 10 in terms of amount of local GC content differences that could be predicted based on the chromatin state annotations there was representation from four different fetal samples and three fibroblast samples (34.5-50.2% predicted) (Table 1). Meanwhile, chromatin state annotations of epigenomes corresponding to brain tissues predicted a lower amount (13.6-28.3%, Supplementary Figure 6, Supplementary Table 2).

**Table 1.** Top epigenomes from Roadmap Epigenomics based on percentage of differences in local GC content between proband and sibling variants predicted by the variants’ chromatin state annotations.

| Roadmap Epigenome ID | Description | Percentage predicted |
| --- | --- | --- |
| E055 | Foreskin Fibroblast Primary Cells skin01 | 50.2 |
| E092 | Fetal Stomach | 48.1 |
| E017 | IMR90 fetal lung fibroblasts Cell Line | 47.1 |
| E090 | Fetal Muscle Leg | 40.6 |
| E118 | HepG2 Hepatocellular Carcinoma Cell Line | 37.3 |
| E080 | Fetal Adrenal Gland | 34.9 |
| E056 | Foreskin Fibroblast Primary Cells skin02 | 34.5 |
| E006 | H1 Derived Mesenchymal Stem Cells | 31.7 |
| E058 | Foreskin Keratinocyte Primary Cells skin03 | 29.9 |
| E034 | Primary T cells from peripheral blood | 29.7 |

For the top 10 epigenomes whose chromatin states collectively predicted the highest percentages of the local GC content differences, we examined the contribution from each individual chromatin state (Methods). We observed that the quiescent state (*Quies*) had the largest contribution to the local GC content differences predicted, with an average of 14.7% across the 10 epigenomes. This state preferentially overlapped sibling variants compared to proband variants in all the 10 epigenomes, with siblings having 9.2% more variants assigned to this state on average (Figure 6, Supplementary Tables 3, 4). We note that the large contribution from the *Quies* state is in part since it is the largest state, overlapping with 42% of all variants on average across the top 10 epigenomes. The state with the second largest contribution on average were the strong transcription state (*Tx*), which constitutes 1.9% of all variants but predicts 4.9% of the local GC content differences and preferentially overlaps proband variants in all the 10 epigenomes (Figure 6, Supplementary Tables 3, 4). The state with the next largest largest contribution on average was the bivalent enhancer state (EnhBiv), constituting 1.1% of the variants while predicting 3.5% of the differences and also preferentially overlapping proband variants in all 10 epigenomes (Figure 6, Supplementary Tables 3, 4).

**Figure 6.**
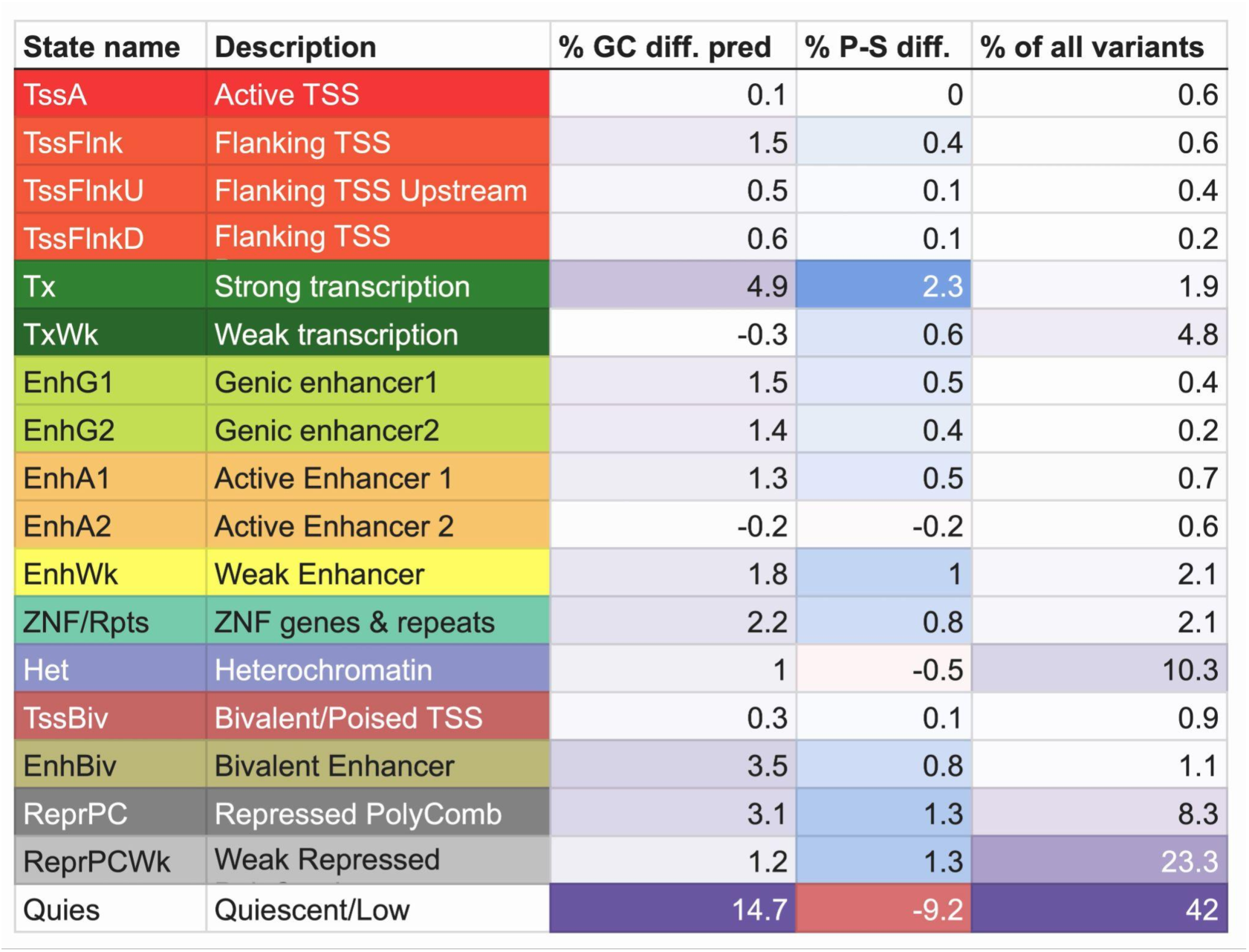
Contributions of each chromatin state to the local GC content rank differences predicted. Percentages shown are mean across the top 10 epigenomes whose chromatin states collectively explained the highest percentages of the local GC content rank differences. % GC diff. pred: mean percentage of proband-sibling local GC content rank difference predicted by the chromatin state. The smallest value is colored white and the largest value is colored purple. % P-S diff.: mean difference between proband and sibling percentages of variants overlapping the chromatin state, with positive indicating preferential overlap with probands. The largest positive value is colored blue and the largest negative value is colored red. % of all variants: mean percentage of variants in the top neighborhoods overlapping each of the states. The smallest value is colored white and the largest value is colored purple.

### Specificity of association signal with respect to sex-upstream/downstream combinations and ASD phenotype

To determine the extent to which the association signal is unique to M-F upstream variants we applied ENSAS to male proband-female sibling downstream (M-F downstream), male proband-male sibling upstream (M-M upstream) and male proband-male sibling downstream (M-M downstream) variants separately. We used the same set-up as the M-F upstream analysis except for the Naive Bayes analysis we we restricted to k-mers of length 6. We observed substantially reduced p-values in these analyses compared to the M-F upstream analysis (Supplementary Figure 7). Using either Bonferroni-based or permutation-based multiple testing corrections we did not observe any significant associations between local GC content and ASD phenotype among the three other combinations after controlling for multiple testing. The most significant p-values for the other combinations were p=8.6×10^-5^ for M-F downstream, 2.1×10^-5^ for M-M upstream, and 2.3×10^-5^ for M-M downstream compared to 1.3×10^-7^ for M-F upstream (Supplementary Figure 7a). The Naive Bayes analysis also did not yield any apparent signal beyond local GC content (Supplementary Figure 7b). Overall these results support the M-F and upstream specificity of the association signal.

To further investigate the specificity of the signal, we asked whether similar male-female differences are seen in an independent dataset unrelated to the ASD phenotype. For this, we applied ENSAS to *de novo* variants from a WGS dataset consisting of 2976 trios of Icelandic origin^31^. After applying the same filtering steps as in the SSC analysis (Methods), we applied ENSAS by comparing all male samples against all female samples and observed no neighborhoods significant for either the local GC content analysis after either Bonferroni-based or permutation-based multiple testing corrections (Supplementary Figure 8a). The Naive Bayes analysis also did not yield any significant findings (Supplementary Figure 8b) This suggests that the identified male-female association signal is not common to all de novo datasets and may be related to ASD phenotypic differences, though we cannot exclude technical differences between the two cohorts.

## Discussion

Here we revisited previous findings of ASD associations based on a deep learning-based approach^11^ and showed that local GC content was sufficient to produce similar ASD associations for noncoding variants excluding CSS variants. We further showed that after conditioning on local GC content the reported significant association signals based on DIS were no longer significant. However, these analyses did not exclude there is additional sequence association signal beyond local GC content. The analyses also did not directly provide insight into why local GC content was sufficient to identify the reported associations including a suggested preferential association for variants differentially expressed in brain tissues. To gain insights into this association we first extended the analyses to consider the sex of the probands and siblings. This suggested among male probands, which represent the large majority of probands, the signal was specific to those that have a female sibling as opposed to a male sibling. We were able to further isolate this signal to variants upstream of their assigned genes.

To more systematically investigate this association signal, we developed ENSAS. ENSAS generalizes the previous analyses by considering sequence signals that can be captured in k-mers instead of only local GC content. In addition, the gene expression neighborhoods it defines are designed to have greater coverage of expression space than the previously defined gene sets based on GTEx differential tissue expression. We applied ENSAS to M-F upstream variants and showed that incorporating k-mer-based sequence context did not lead to overall improved prediction of proband-sibling status compared to local GC content given current sample sizes. However, there still exists the possibility that sequence information beyond local GC content could identify enhanced association signals. The top neighborhoods where proband-sibling local GC content differ the most showed the strongest enrichment for synapse-related GO terms, consistent with previous findings in ASD *de novo* coding variants^6,28^. We identified epigenomes whose chromatin states could collectively predict a large portion of the proband-sibling difference in the top neighborhood, primarily those pertaining to fetal and fibroblast samples. Across these epigenomes we observed the largest contribution to the GC content predicted based on chromatin state annotations from the preferential overlap of the quiescent state among sibling variants and the preferential overlap of the strong transcription state and bivalent enhancer state among proband variants the latter of which has a strong association with developmental functions^32,33^.

Understanding why the signal was specific to male probands with female siblings could be an avenue of investigation for future studies. One hypothesis to investigate is that some of the mutations occurred post-zygotically early enough in development^34^ to be found in a large fraction of cells and sex specific chromatin or transcriptomic differences could be associated with the mutational differences^35^. Also understanding why the observed signal is specific to variants in regions upstream of TSS relative to downstream could also be an avenue for future investigation, but could be related to different transcriptional processes or chromatin environments associated with these regions. We did show the difference cannot be explained by variants in the immediate upstream promoter regions, for which a previous association with ASD^13^ was reported.

We note that here ENSAS was only applied to *de novo* variants. A possible area of future investigation is to apply ENSAS to rare variants from non-family-based case-control studies. However, this can be a challenging task, and effective application may require incorporation of stringent variant filtering thresholds such as variants not already present in aggregation databases^36^. In addition, ENSAS does not directly determine if sequence differences are due to biological differences or technical confounders associated with sequencing^19^. We note that following Zhou *et al.*^11^, we used a repeat-masked subset of variants and thus have excluded a set of variants more vulnerable to technical confounders^18^. In addition we showed that the association signal appears mainly driven by proband-sibling pairs with matching sequencing lane information thus suggesting sequencing differences is less likely an explanation. The specificity of the signal particularly for brain associated genes, male probands with female siblings, and variants upstream of genes relative to downstream are also suggestive of a biological basis as it would exclude technical confounders that would not display this level of specificity. We note that even assuming the association signal is not driven by technical sequence confounders we cannot exclude the possibility that there is a biological basis for it that is correlated with the phenotype but is not directly causally related.

The role of rare noncoding variants in complex phenotypes remains an underexplored area. With the increasing availability of large-scale WGS datasets, there is a need for effective analytical frameworks. Here we presented an analytical framework that simplifies and extends previous results on the role of non-coding de novo variants in ASD and holds promise for its application in other WGS studies.

## Methods

### Defining local GC content

For each of the 127,140 *de novo* variants from Zhou *et al.*^11^, which were all autosomal single nucleotide variants not overlapping repetitive elements, we extracted the DNA sequence that was within 100 bp on each side to have sequences of length 201bp using the GRCh37 assembly from UCSC genome browser^37^ release 14. We then counted the number of nucleotides that were a ‘G’ or ‘C’ in the 201bp for each sequence.

### DIS adjusted for local GC content

To construct a DNA DIS score adjusted for local GC content, for each variant we took the mean DNA DIS score of all other variants that had the exact same number of ‘G’ or ‘C’ nucleotides within 100bp and then subtracted that value from the variant’s DNA DIS value. We note that the DIS values globally were already mean-centered around 0. We repeated the same procedure for the RNA DIS, except restricted to the 77,157 variants that had an RNA DIS available. There were four variants for the DNA DIS that had a unique count for the number of ‘G’ or ‘C’, which were outliers, and for these variants we did not subtract any value, and similarly for three outlier variants for the RNA DIS.

### Defining coding regions and canonical splice sites

Throughout our analyses, coding regions are defined by Gencode^38^ v24 annotations lifted to GRCh37, and canonical splice sites (CSS) are defined by any variants annotated as a ‘splice_acceptor_variant’ or ‘splice_donor_variant’ by the Ensembl Variant Effect Predictor (VEP)^39^ release 109.3 for GRCh37 release 98.

### Genomic variant set analysis

For computing the significance of proband and sibling score differences for different sets of genomic variants, we followed the same procedures as used by Zhou *et al.*^11^ and extended their code for the version of the analysis that excluded coding variants. We used the version of the variant annotations used by Zhou *et al.*^11^, which was provided by them. We note there are small differences between that version and the version of the annotations provided in Supplementary Table 1 of Zhou *et al.*^11^ for the RNA gene sets, due to differences in the gene identifiers used. For filtering variants in coding regions we used Gencode v24^38^ lifted to GRCh37.

We tested the 130 variant sets using: (1) local GC content, (2) the original DNA and RNA DIS reported (60 sets were tested with DNA DIS, 70 sets were tested with RNA DIS) (3) the DNA and RNA DIS adjusted for local GC content, described above. The p-value significance of proband mutations was computed with a one-sided one-sided Mann-Whitney U-test and false discovery rates were computed with the Benjamini–Hochberg procedure. We also conducted additional sets of tests for each score where we first removed any variant in coding regions or CSS from the variant set and then applied the same procedures.

### Tissue-specific gene expression analysis

We analyzed tissue-specific gene expression from GTEx following the procedures of Zhou *et al.*^11^ Each of the variants was assigned to their respective genes based on the distance to a nearest representative TSS as described by Zhou *et al*.^11^ We used the list of GTEx tissue-specific genes for each of the 53 GTEx tissues provided by Zhou *et al.*^11^, which was defined as those with five times its median expression across all tissues. We note that for this analysis the authors created a single combined score of the RNA and DNA DIS. We followed the author’s procedure in their provided code for creating the combined score. Specifically in this analysis, if a variant had an RNA DIS value available the mean of the DNA and RNA DIS was used, otherwise just the DNA DIS was used. Also following the code provided by the authors, only variants within 100kbp of a TSS or intron variants within 400bp of an exon boundary were included.

For each of the GTEx tissues, we performed a one-sided Mann-Whitney U-test on the local GC content of proband variants versus sibling variants assigned to these genes. We repeated the same test using DIS scores and DIS scores corrected for local GC content, and with coding and CSS variants removed. The multiple testing correction was done with the Benjamini–Hochberg procedure.

To separately test variants upstream of the outermost TSS of their assigned gene from those downstream we remapped the assignment of variants to genes. For each gene we determined the position of the outermost TSS, which was the lowest coordinate if the gene was on the positive strand and the greatest coordinate if the gene was on the negative strand. We used the same set of gene annotations used for defining coding regions and CSS as described above.

We then assigned each variant to its nearest outermost TSS. Variants with a coordinate less than the TSS of its assigned gene where the gene is on the positive strand or with a coordinate greater than the TSS of its assigned where the gene is on the negative strand were classified as upstream variants. The remaining variants were classified as downstream variants.

We then repeated the above testing procedure separately using different subsets of variants: (A) Variants from proband-sibling pairs where the proband is male and the sibling is female. (B) Variants from proband-sibling pairs where the proband is male and the sibling is male. (C) Same as (A) but the variant is within 100kbp upstream of the nearest TSS (D) Same as (A) but the variant is within 100kbp downstream of the nearest TSS For (A), (B), (C), and (D) we removed coding and CSS variants. For (C) and (D) we did not include variants based on proximity to exon boundaries.

### Defining gene expression neighborhoods

To define gene expression neighborhoods we first obtained the pairwise gene-gene co-expression correlation matrix from Geneshot^22^, which is computed using the RNA-seq data compiled by the ARCHS4 resource^23^. The matrix contains a total of 29,820 genes. We defined the “neighborhood” *N_g_* surrounding each gene *g* to be the top *M* variants that are assigned to its closest genes based on the pairwise distances, including the variants assigned to *g* itself, with ties broken arbitrarily. This strategy is similar to the gene set augmentation functionality of Geneshot^22^. We also greedily pruned variants that are <*L*/2bp apart, that is for each pair of variants with <*L*/2bp between each other we kept the variant with smaller coordinates.

### ENSAS: Sequence analysis of the gene expression neighborhoods

To test for the local GC content differences between two groups of variants in a neighborhood, ENSAS uses a Mann-Whitney U-test which could be either one-sided or two-sided depending on the specific application. ENSAS computes a Bonferroni-corrected p-value threshold of 0.05 / *n* where *n* is the number of neighborhoods.

To test for the higher-order sequence differences between the two groups, ENSAS uses a machine-learning-based approach. For each variant in a neighborhood ENSAS extracts the (*L*+1)bp nucleotide sequence centered around it. Then for a given *k*, it counts the number of occurrences of each k-mer in each variant. For each neighborhood, the variants are split into evenly sized training and testing folds. A multinomial Naive Bayes classifier is trained on the training fold to distinguish between variants in the two groups, using each variant’s k-mer occurrence counts as features and uniform class priors. One of the groups is arbitrarily labeled as positive and the other group is labeled as negative. The classifier is then applied to the testing fold to compute the posterior probability (“score”) of each testing variant being in the positive group. ENSAS then tests the difference in the distribution of scores between the two groups of variants in the testing with a one-sided Mann-Whitney U-test. The same test is also performed using local GC content on the testing fold. ENSAS by default computes a Bonferroni-corrected p-value threshold of 0.05 / n where n is the number of neighborhoods. As described below ENSAS, also provides a permutation based multiple testing correction option.

We applied ENSAS to variants upstream of TSS in male proband-female sibling pairs. Given the high correlation of local GC content and DIS, for this application we used one-sided tests for local GC content differences to maintain consistency with Zhou et al.^11^’s use of one-sided tests for DIS. Across the GTEx tissues, the mean number of variants (within the male proband-female sibling upstream subset) whose genes show differential expression is 1048. We therefore set the size of neighborhoods *M* to be 1000 to approximate that. We set the sequence length *L* to be 201. We separately tested 1,2,3,4,5,6 and 7-mers. For the other sex-upstream/downstream combinations (male proband-female sibling downstream, male proband-male sibling upstream, male proband-male sibling downstream) we also performed ENSAS with neighborhood sizes of 1000 and a k-mer length of 6.

### Permutation-based multiple testing correction

In addition to the Bonferroni-based correction, ENSAS also provides a permutation-based approach to correct for multiple testing. Specifically, in each permutation ENSAS randomly swaps the proband-sibling labels within each pair. Let the Mann-Whitney U p-value of a neighborhood *i* on real data be *p_i_*. The number of discoveries claimed as true positives based on threshold *p_i_* is then *|{j: p_j_ < p_i_}|* that is the number of neighborhoods with a smaller p-value. The estimated number of false positive discoveries based on *B* (1000 by default) permutations is Σ *|{j: p^b^ < p}|* / B that is the average number of neighborhoods with smaller p-value than *p*_i_ across the *B* permutations, where *p_j_^b^* is the p-value of neighborhood *j* in the *b*^th^ permutation. The estimated FDR based on threshold *p_i_*is the estimated number of false positive discoveries divided by the number of discoveries claimed as true positives. ENSAS finds *i* such that the estimated FDR is 0.05 and declares all neighborhoods with p-value smaller than *p_i_* as significant.

### Gene ontology enrichment analysis for the top neighborhoods

We performed Gene Ontology (GO) enrichment analyses for the top neighborhoods with the most significant local GC content p-values from the ENSAS performed on M-F upstream variants. For the GO analyses, we used as the foreground the set of genes with assigned variants in each neighborhood and separately two different sets of background genes: (1) union of all genes assigned to the M-F upstream variants and (2) the union of genes assigned to the M-F upstream variants that were also considered differentially expressed within any of the 13 brain-related GTEx tissues^11^. The analysis was done using GOATOOLS^40^, with a Fisher’s exact test conducted for each GO term and FDR-controlling Benjamini-Hochberg procedure to correct for multiple testing.

### Evaluating ENSAS with Simulations

We investigated if ENSAS can differentiate between proband and sibling variants simulated to have a different underlying chromatin state distribution. To do this we used chromatin state annotations produced by a 18-state ChromHMM model from the Roadmap Epigenomics Consortium^29,30^. We selected an epigenome (E003-H1 cell line) and simulated 500 proband and 500 sibling labeled variants by randomly drawing positions (excluding assembly gaps) belonging to different chromatin states. Among proband variants, X% were drawn from the four active TSS-associated states (*TssA*, *TssFlnk*, *TssFlnkU* and *TssFlnkD*) and the other (100-X)% were uniformly drawn across the genome, where X was a parameter we varied. All sibling variants were uniformly drawn across the genome. We greedily pruned variants such that no two variants were <100bp apart to ensure no overlap of sequences. For each X in {10, 20, 50, 80, 100} we generated 50 simulated datasets. We applied ENSAS to distinguish between the positive and negative variants, with a sequence length L of 201bp and a neighborhood size M of 1000. We used 1-7 as candidate lengths of k-mers and repeated the train-test split 50 times.

### Prediction of proband-sibling sequence differences in the top neighborhood with chromatin states

For the top neighborhood identified when applying ENSAS to M-F upstream variants, we quantified the extent by which chromatin state annotations in particular epigenomes can predict the proband-sibling differences in the local GC content. We annotated each of the 1000 variants using the 18-state ChromHMM model defined based on six histone modifications across 98-reference epigenomes by the Roadmap Epigenomics Consortium^29,30^. For each variant, we converted its local GC content to a rank value and broke ties arbitrarily, as the significance tests ENSAS applied were rank-based. We then randomly partitioned the variants into five subsets. For each variant, we predicted its rank based on the mean rank of variants from the other four subsets than which the variant belonged. For this prediction we used variants assigned to the same chromatin state regardless of whether the variant was from a proband or sibling. If the state is not present in the training set then the mean rank of all variants is assigned as the prediction. We took the difference between the proband mean predictions and the sibling mean predictions and divided by the observed mean difference in rank to determine the percent of rank differences predicted by the chromatin state annotations in a reference epigenome.

To quantify the contribution of individual chromatin states to the mean prediction rank differences between proband and siblings we computed the quantity ((*a_proband,s_*-*r*)\**f_proband,s_* - (*a_sibling,s_*-*r*)\**f_sibling,s_*). *a_proband,s_* and *a_sibling,s_* are the mean rank predictions for variants in state *s* from proband and siblings, respectively. *f_proband,s_* and *f_sibling,s_* are the fraction of predictions for variants in state *s* from proband and siblings, respectively. *r* is the overall mean rank of all variants. We then divided this quantity by the observed mean difference in rank and multiplied this value by 100%. We note that this value can be negative for some states.

### Stratification by matching sequencing lanes

We obtained the sequencing lane information for each sample, represented by columns “LANE_from_ID” and “LANE_from_PU” in Supplementary File 1. We defined proband-sibling pairs with matching sequencing lanes to be the pairs where the proband and the sibling samples have exactly the same lane assignments in both columns, and vice versa for mismatched lanes. Under this criteria there were 968 samples from male proband-female sibling pairs with matching sequencing lanes and 606 samples from those pairs with mismatched sequencing lanes. Next, for each neighborhood in the ENSAS on the M-F upstream variants, we performed proband vs. sibling Mann-Whitney U-tests separately on samples with matching sequencing lanes and samples with mismatched sequencing lanes.

### Removing variants in promoter regions

From the 12,293 M-F upstream variants, we removed variants in promoter regions, defined as <2kbp upstream of their assigned nearest outermost TSS following the promoter definition from An *et al*.^13^ We then performed ENSAS on the remaining 11,395 M-F upstream variants with neighborhood size of 1000.

### Application to an Icelandic dataset

We applied ENSAS to a dataset of an Icelandic population consisting of 194,687 autosomal *de novo* variants from 2976 WGS trios reported by Halldorsson *et al.*^31^ We lifted the variants to GRCh37 and performed comparison between male samples and female samples. For the local GC content analysis we tested whether variants from male samples had higher local GC content than those from female samples using a one-sided test. Using the same procedure that was applied to the SSC variants, we removed indels, removed variants in coding regions and CSS, only kept variants <100kbp upstream of their nearest outermost TSS, and greedily pruned variants that are <100bp apart. We applied ENSAS with 1000 neighbors to the remaining 41,846 variants.

## Data availability

*De novo* variant calls from the SSC WGS data are obtained from Ref^11^. *De novo* variant calls from the WGS data for the Icelandic population are obtained from Ref^31^. Gene definitions are obtained from Gencode v24 lifted to GRCh37: https://www.gencodegenes.org/human/release_24lift37.html. Ensembl release 109.3 variant annotations are obtained from https://github.com/Ensembl/ensembl-vep/tree/release/109.3. Geneshot gene expression correlation matrix is obtained from https://maayanlab.cloud/geneshot/download.html. 18-state ChromHMM annotations for 98 epigenomes are obtained from the Roadmap Epigenomics Project: https://egg2.wustl.edu/roadmap/data/byFileType/chromhmmSegmentations/ChmmModels/core_K27ac/jointModel/final/

## Code availability

Source code for ENSAS can be found at https://github.com/ernstlab/ENSAS

## Acknowledgements

We acknowledge the contributions of participants in the SSC and Iceland cohorts. We thank Jian Zhou, Christopher Park, Chandra Theesfeld, and Olga Troyanskaya for discussions and sharing code and data. We thank Daniel Geschwind and Stephan Sanders for discussions and comments on a preliminary draft. We thank Shan Dong and Stephan Sanders for providing annotations of the inferred sex of individuals in SSC data. We thank Bjarni V. Halldórsson for providing annotations of the inferred sex of individuals in the Iceland cohort. We thank members of the Ernst lab and whole-genome sequencing for psychiatric disorders (WGSPD) consortium for discussions. We acknowledge funding from US National Institutes of Health grants DP1DA044371, U01MH105578, R01MH109912, R01MH110927, U01HG012079, U01MH130995, and UCLA Jonsson Comprehensive Cancer Center and Eli and Edythe Broad Center of Regenerative Medicine and Stem Cell Research Ablon Scholars Program.

## Author Contributions

R.L and J.E conceived & developed the methods, performed analyses and wrote the manuscript.

