## Supplementary information for "Identifying associations of *de novo* noncoding variants with autism through integration of gene expression, sequence and sex information"

**Runjia Li[1] and Jason Ernst[1,2,3,4,5,6,7]**
[1]Bioinformatics Interdepartmental Program, University of California, Los Angeles, CA, USA.
[2]Department of Biological Chemistry, University of California, Los Angeles, CA, USA.
[3]Eli and Edythe Broad Center of Regenerative Medicine and Stem Cell Research at University of California, Los Angeles, CA, USA.
[4]Computer Science Department, University of California, Los Angeles, CA, USA.
[5]Jonsson Comprehensive Cancer Center, University of California, Los Angeles, CA, USA.
[6]Molecular Biology Institute, University of California, Los Angeles, CA, USA.
[7]Department of Computational Medicine, University of California, Los Angeles, CA, USA.

### Supplementary figures

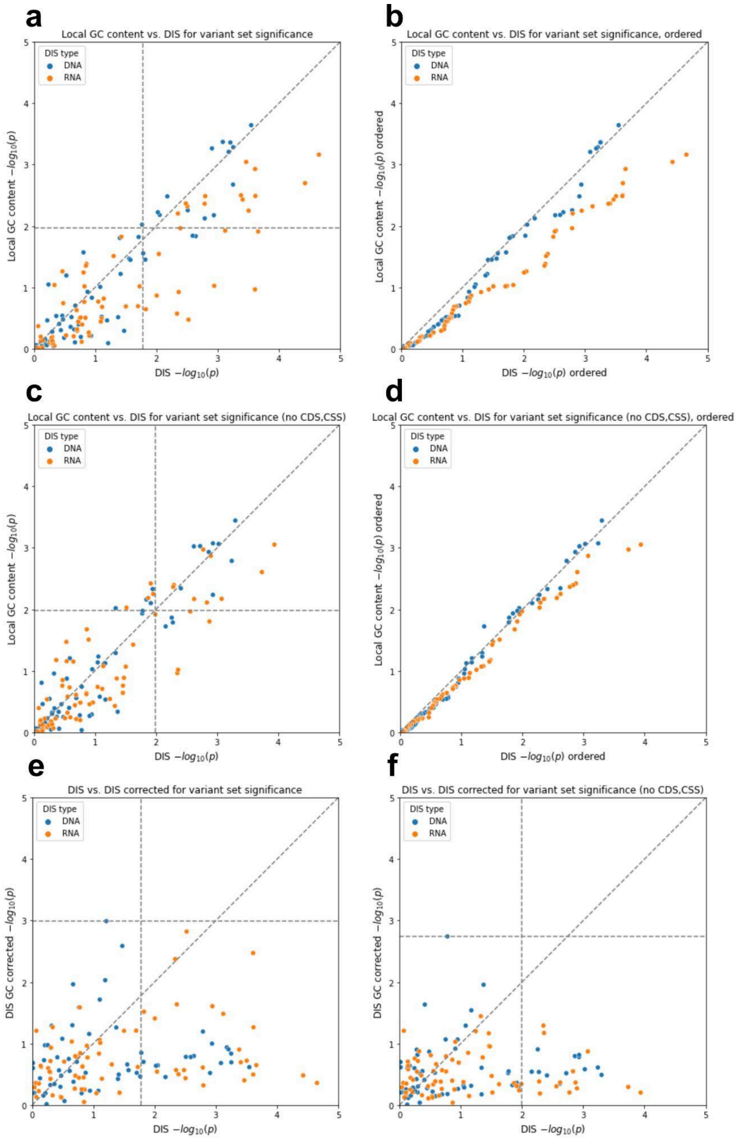


**Supplementary Figure 1** -log10 p-values from the genomic variant set analysis using the same testing procedure as described previously[1]. All analyses were restricted to noncoding variants. Both the x- and y-axis show -log10 p-values from one-sided Mann-Whitney U-tests. Horizontal and vertical dashed lines show p-values at FDR threshold of 0.05. Points greater than (but not on) these lines are significant after FDR correction. Diagonal dashed lines show the unit slope. **(a)** proband-sibling differences for local GC content vs. proband-sibling differences for DIS with DNA and RNA-based tests colored separately **(b)** same as (a) but the scatter plot shows the *i*th most significant p-value for DIS against local GC content separately for the DNA and RNA based tests for all possible values of *i*. **(c)** same as (a) but with CSS variants removed; **(d)** same as (b) but with CSS variants removed **(e)** proband-sibling differences for DIS (corrected for local GC content) vs. proband-sibling differences for uncorrected DIS; **(f)** same as (e) but with CSS variants removed.

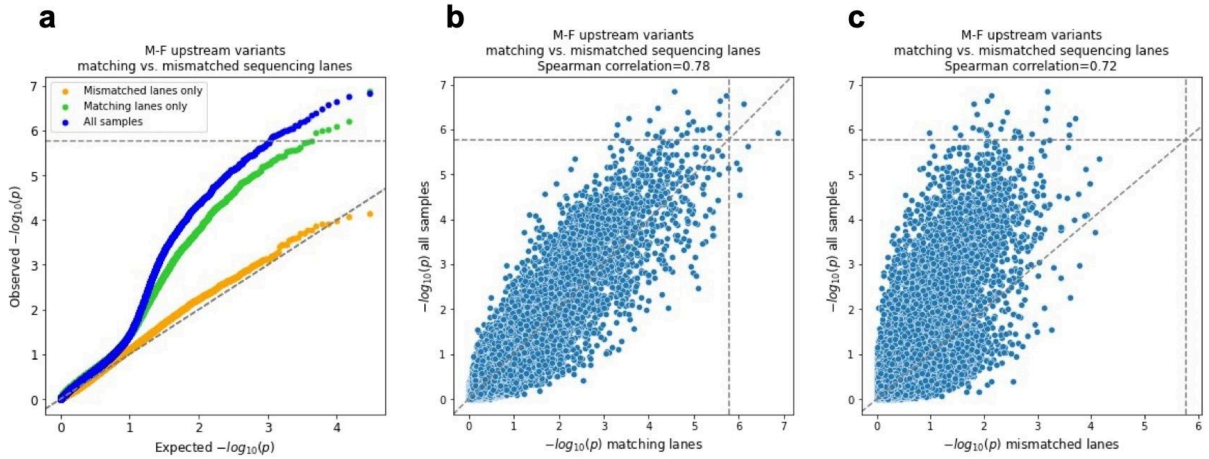


**Supplementary Figure 2** Comparison of local GC content Mann-Whitney U p-values for ENSAS on M-F upstream variants of all samples, samples with matching sequencing lanes with their probands/siblings (“matching lanes”), and samples with mismatched sequencing lanes with their probands/siblings (“mismatched lanes”). Horizontal and vertical dashed lines show Bonferroni-based p-value multiple testing significance thresholds of 0.05 / n where n=29,820 is the number of neighborhoods. Diagonal dashed lines show the unit slope. **(a)** QQ-plots for the neighborhood local GC content p-values of three sample groups. **(b)** Neighborhood p-values for all samples vs. samples with matching sequencing lanes. **(c)** Neighborhood p-values for all samples vs. samples with mismatched sequencing lanes.

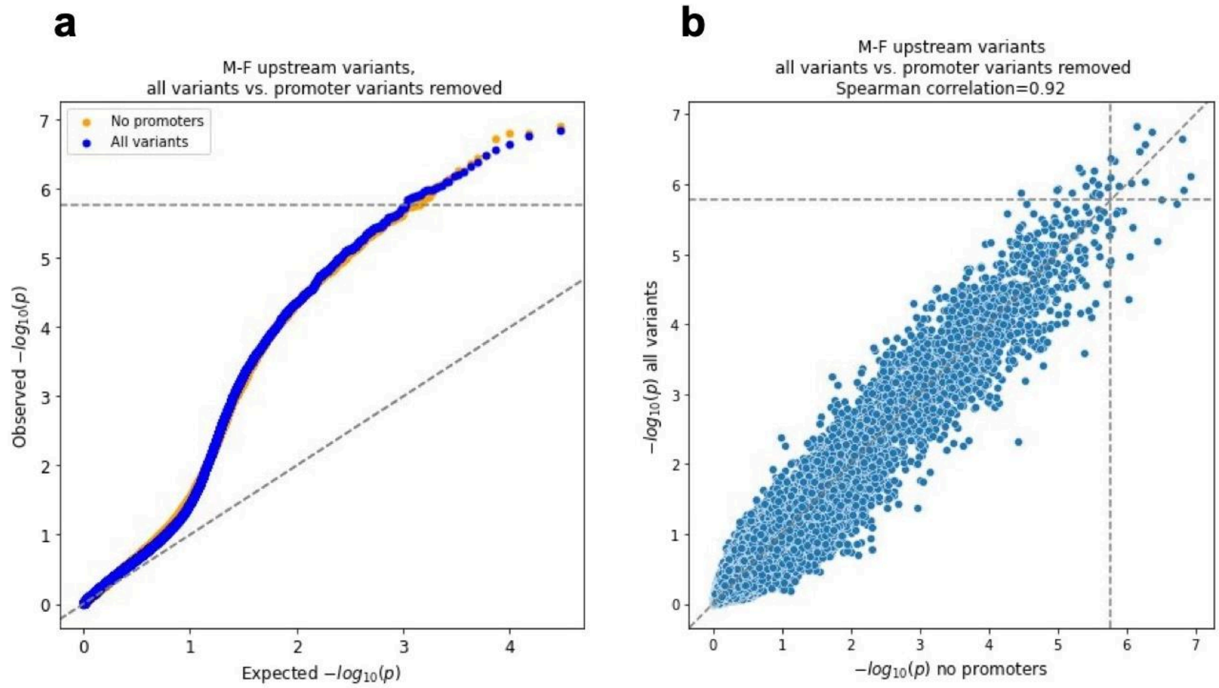


**Supplementary Figure 3** Comparison of local GC content Mann-Whitney U p-values ENSAS on all M-F upstream variants vs. those excluding promoter variants. Horizontal and vertical dashed lines show Bonferroni-based p-value multiple testing significance thresholds of 0.05 / n where n=29,820 is the number of neighborhoods. Diagonal dashed lines show the unit slope. **(a)** QQ-plot of neighborhood p-values before and after removing promoter variants. **(b)** Neighborhood p-values for all variants vs. promoter variants removed.

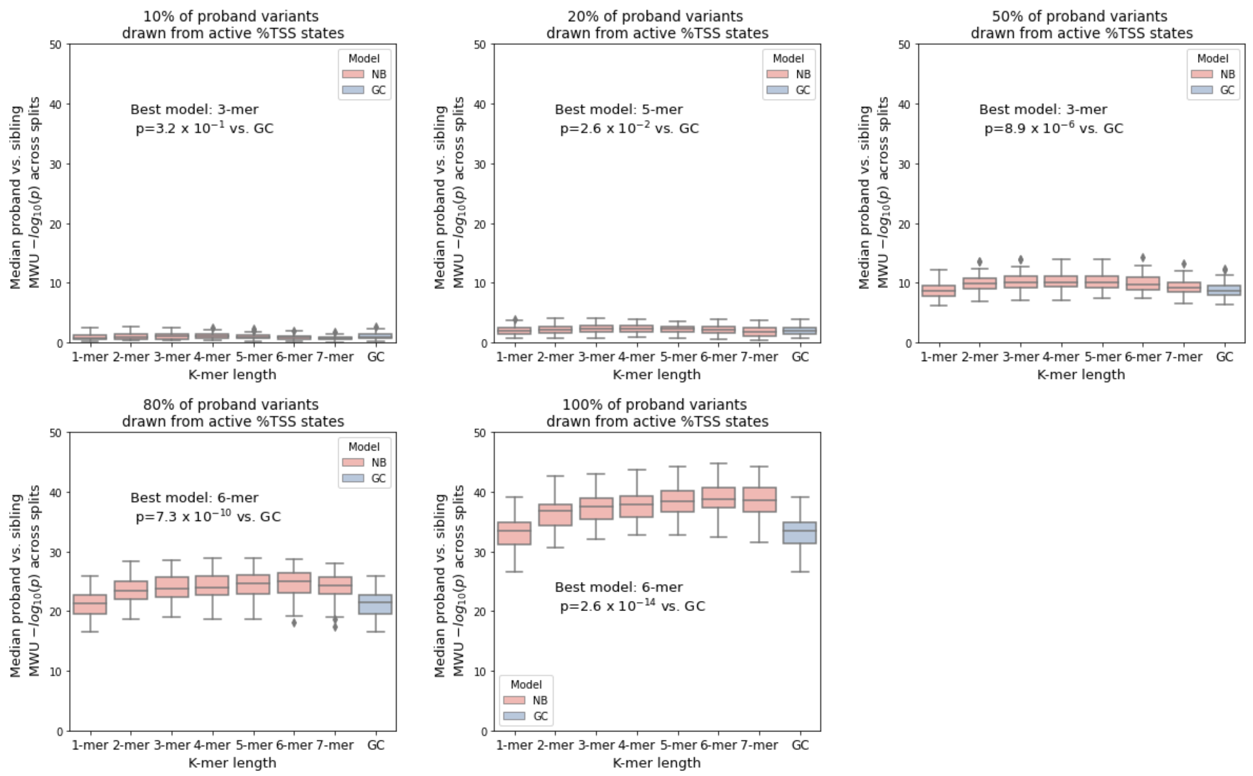


**Supplementary Figure 4** Median Naive Bayes (NB) model and local GC content (GC) Mann-Whitney U p-values of proband vs. sibling variants across 50 random train-test splits on simulated datasets. Each subfigure represents 50 simulated datasets where a fixed portion of proband variants were drawn from active TSS states (TssA, TssFlnk, TssFlnkU and TssFlnkD, Methods). Boxes show medians and interquartile ranges across 50 simulated datasets. Points above or below 1.5x interquartile range are drawn as outliers. Annotated text shows the best-performing k-mer model in terms of median performance across simulations and the one-sided Mann-Whitney U p-values of its performance vs. local GC content.

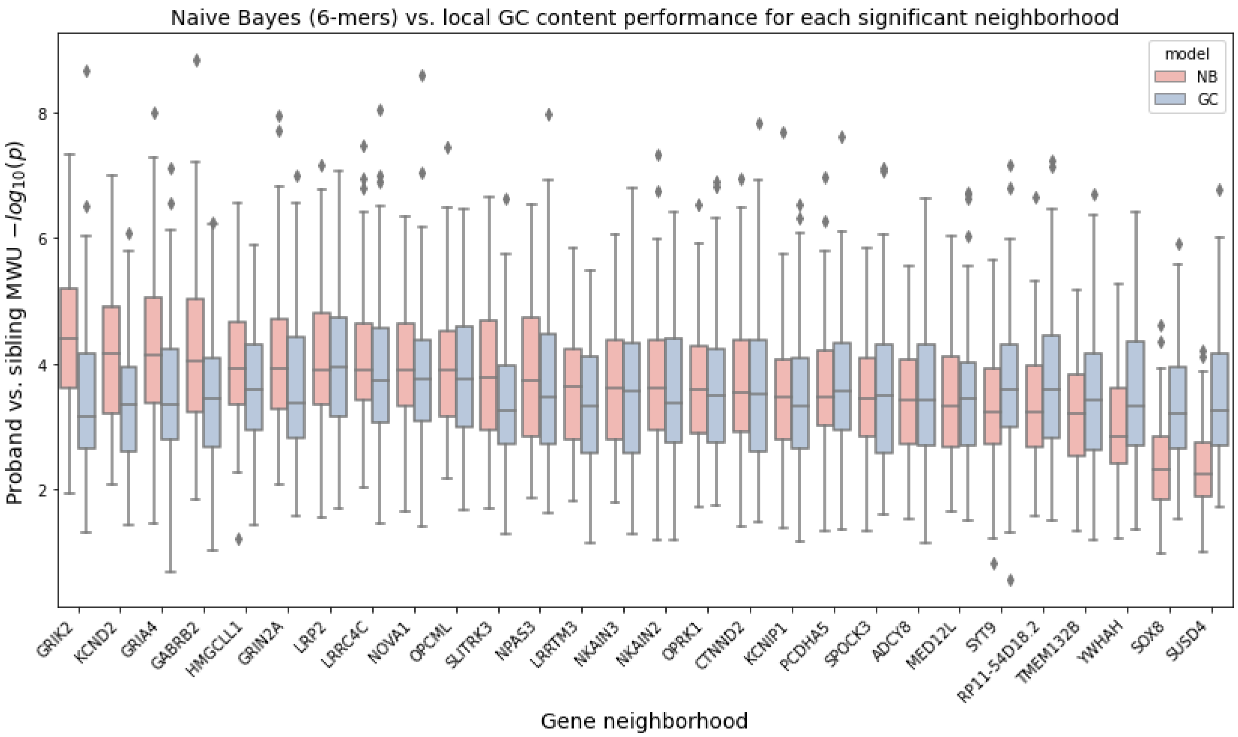


**Supplementary Figure 5** 6-mer Naive Bayes model (NB) and local GC content (GC) Mann-Whitney U p-values across 100 random train-test splits for each of the 28 significant neighborhoods. Boxes show median and interquartile ranges. Points above or below 1.5x interquartile range are drawn as outliers. The neighborhoods are sorted by their median Naive Bayes p-values.

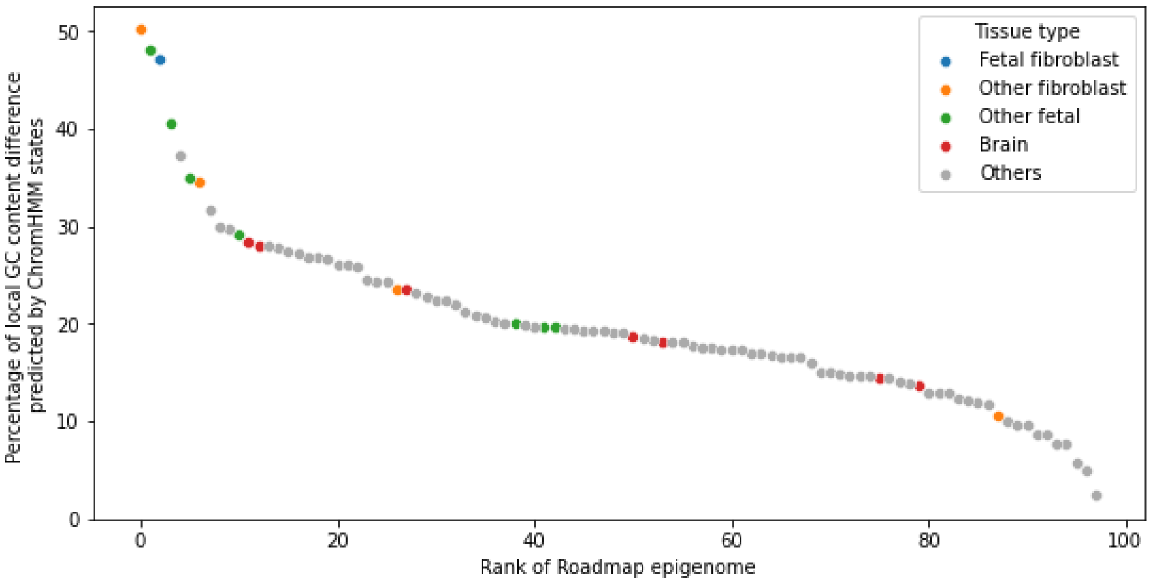


**Supplementary Figure 6** Percentage of differences in local GC content between proband and sibling variants predicted by the variants' chromatin states, ranked for each of the 98 epigenomes from Roadmap Epigenomics[2]. Selected subsets groups of epigenomes are colored as follows: Blue: fetal fibroblast tissues; Orange: non-fetal fibroblast tissues; Green: non-fibroblast fetal tissues; Red: brain tissues; Grey: all other tissues.

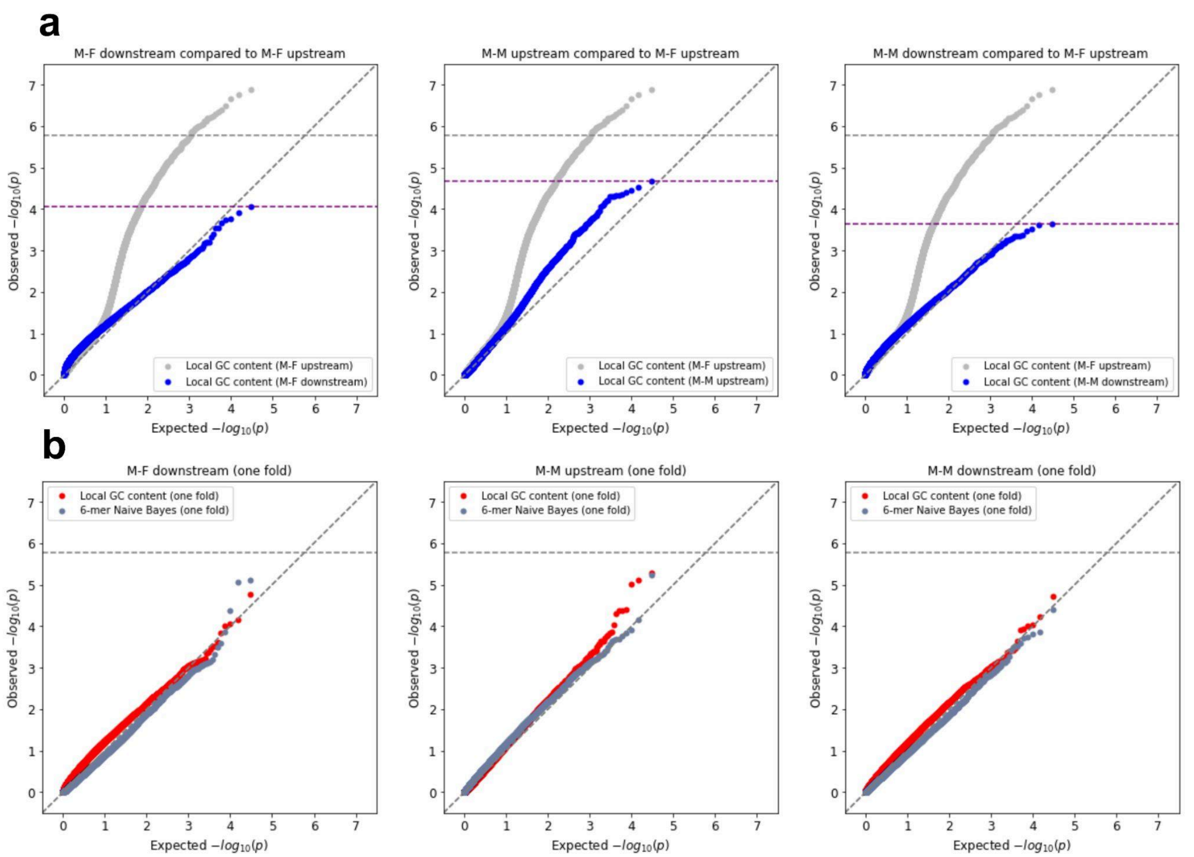


**Supplementary Figure 7** QQ-plots for Mann-Whitney U p-values for ENSAS on M-F downstream (left), M-M upstream (middle), and M-M downstream variants (right). Grey horizontal dashed line shows Bonferroni-based p-value multiple testing significance threshold of 0.05 / n where n=29,820 is the number of neighborhoods. Purple horizontal dashed lines show permutation-based multiple testing threshold at an FDR of 0.05, with points greater than (but not on) these lines significant after correction. Diagonal dashed lines show the unit slope. **(a)** P-values for local GC content using all variants in the neighborhood are shown. P-values for the SSC M-F upstream analysis are also displayed for comparison. **(b)** P-values for local GC content and 6-mer Naive Bayes model on the testing fold for each neighborhood are shown.

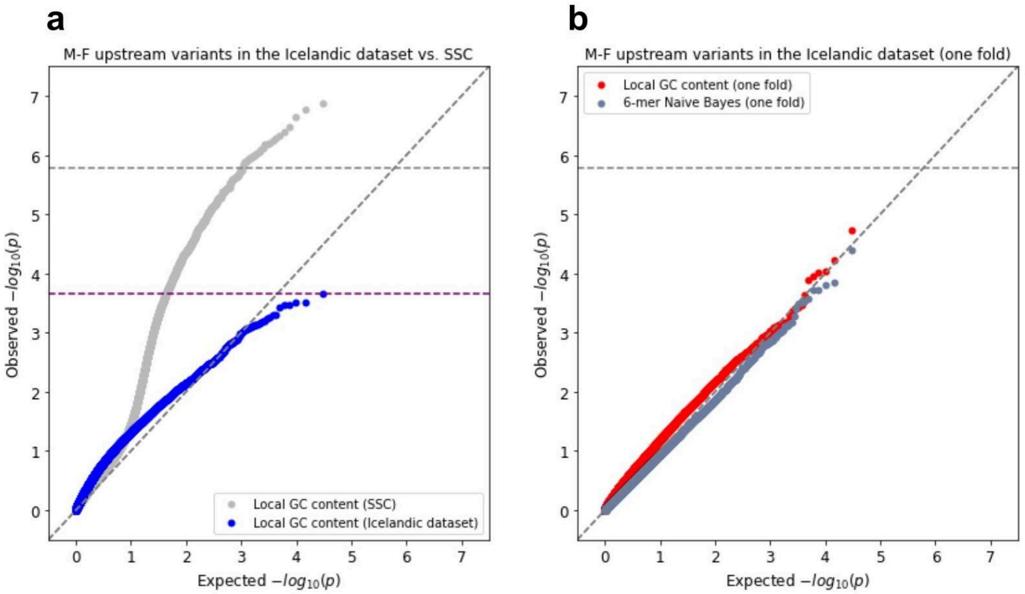


**Supplementary Figure 8** QQ-plot of Mann-Whitney U p-values for ENSAS on all M-F upstream variants in the Icelandic dataset[3]. Grey horizontal dashed line shows the Bonferroni-based p-value multiple testing significance threshold of 0.05 / n where n=29,820 is the number of neighborhoods. Purple horizontal dashed line shows permutation-based multiple testing threshold at an FDR of 0.05 (Methods), with points greater than (but not on) these lines significant after correction. Diagonal dashed line shows the unit slope. **(a)** P-values for local GC content using all variants in the neighborhood are shown. P-values for the SSC M-F upstream analysis are also displayed for comparison. **(b)** P-values for local GC content and 6-mer Naive Bayes model on the testing fold for each neighborhood are shown.
